# NK cell-macrophage interactions in granulomas correlate with limited tuberculosis pathology

**DOI:** 10.1101/2025.02.14.638234

**Authors:** Paula Niewold, Marieke E IJsselsteijn, Karin Dijkman, Noel de Miranda, Tom H M Ottenhoff, Frank A W Verreck, Simone A Joosten

**Author notes:** Corresponding author: Simone A Joosten, Albinusdreef 2, C-05-043, 2333 ZA Leiden. The authors have declared no conflict of interest.

## Abstract

Development of novel vaccines and treatment approaches against tuberculosis are hampered by limited knowledge of what constitutes a protective immune response against *Mycobacterium tuberculosis* (*Mtb*). Granulomas are organized immune aggregates that form in the lung in response to mycobacterial infection and are an important site of pathogen-host interaction. The composition and cellular microenvironment within the granuloma impacts the bacterial control capacity. To identify protective responses in granulomas, imaging mass cytometry was used to study archived lung tissue from low dose *Mtb*-infected non-human primates presenting with various levels of disease. This approach revealed that granuloma composition is correlated with the severity of lung pathology. Granulomas of animals with limited lung pathology were enriched for NK cells showing increased interactions with tissue macrophages. This work improves our understanding of local immune interactions in the lung and how these correlate with severity of tuberculosis disease.

**Author summary:** Tuberculosis remains the deadliest infectious disease in the world with 10 million new infections and 1.5 million deaths annually. The existing vaccine is not effective enough to halt the tuberculosis epidemic. In order to develop an improved vaccine, more complete and detailed knowledge on the immune responses that can successfully protect against tuberculosis is required. Individuals that are exposed to *Mycobacterium tuberculosis*, the pathogen that causes tuberculosis disease, fall along a spectrum, with some individuals showing a strong innate capacity to protect against disease, while others fall ill. By studying these differences in a model organism, namely non-human primates, we can learn about protective immune responses. In primates with limited disease, the immune cell aggregates surrounding the bacteria in the lung contained more NK cells than were found in severe disease. In addition, interactions between NK cells and macrophages, cells that can eat bacteria, were seen a lot more in protected animals. This suggests that the interaction between NK cell and macrophages contributes to controlling tuberculosis disease, this knowledge can contribute to improved vaccine strategies.

## Introduction

Tuberculosis remains the deadliest infectious disease in the world and has significant impact particularly on the global south (1). Treatment generally consists of several months of multiple antibiotics, afflicting therapy adherence due to serious side effects, while its efficacy is undermined by rapidly increasing antibiotic resistance (1, 2). Despite extensive efforts to develop novel preventive TB vaccines, the Bacille Calmette Guerin (BCG) vaccine with its variable and limited efficacy in adolescents and adults remains the only licensed vaccine to date (3). Expanding our incomplete knowledge of what constitutes a protective response against *Mycobacterium tuberculosis* (*Mtb*) is essential to the development of improved vaccines and therapeutic options.

The current understanding of protective immunity against pulmonary TB in man is largely based on peripheral blood analysis correlated to disease status, which exists along an imperfectly defined spectrum (e.g. healthy control, house-hold contact, resister, infected, active disease) (4). In addition, it is unclear how reflective these peripheral correlates are for the immune events occurring at the primary site of infection. During Mtb infection, immune cells are attracted to the lung and form structures called granulomas which are the pathological hallmark of TB, but also synapses of pathogen-host interaction playing a key role in mycobactericidal control (5). Granulomas generally consist of fibroblasts, epithelial cells, granulocytes, macrophages, lymphocytes and their organization and composition impacts function (6–10). In the past, studying (granuloma) tissue implied choosing between low dimensional techniques preserving spatial context and higher dimensional analyses on digested tissue (11). Recent technological developments have enabled high dimensional spatial analysis, yielding novel insights through studying cell interactions to understand outcomes of disease i.e. in cancer (12), Sars-Cov2 (13) and inflammatory bowel disease (14). Such technologies have been applied to human TB specimens, showing a highly immune-modulatory environment created by *Mtb* (15) and granuloma heterogeneity (16). While informative, in patients it is typically impossible to determine the exact time and dose of infection, and scarcely available lung material is often limited to late stage disease resections. Therefore, fundamental questions are addressed in controlled *Mtb* infection models like non-human primates, which recapitulate human immunology and TB disease development well (17).

To understand early events in the granulomatous response and its possible impact on later stage disease we used archived lung material of non-human primates collected 6 weeks post IGRA conversion following low-dose infection with *Mtb* (18). Imaging mass cytometry, which accomplishes resolution high-dimensional imaging at 1 µm2 spatial resolution with metal-labelled antibodies (19), was applied to determine cellular composition and spatial organization of 28 granulomas across 6 animals. Analysis revealed granuloma composition and lung pathology were correlated. Specifically, NK cell numbers were increased in granulomas of animals with limited lung pathology and these cells showed enhanced interactions with tissue macrophages. Overall, we mapped cellular phenotypes and interactions within the granulomas to better understand how local interactions in the lung are associated with the extent of pathology.

## Results

To investigate granuloma composition, we used banked formalin-fixed paraffin-embedded (FFPE) lung tissue from macaques that were endobronchially infected with a low-dose of *Mtb* Erdman and sacrificed 6 weeks post interferon gamma release assay (IGRA) conversion. Six macaques representing the spectrum of pathology scores and *Mycobacterium tuberculosis (Mtb)* tissue burden in the endobronchially targeted lung lobe were selected. Hematoxylin & eosin staining was performed on the lung tissue of all 6 animals to visualize the architecture of the lung tissue and location of granulomas (Figure 1C). In each section, 5 granulomas of varying sizes (< 0,3 mm^2^, 0,3 – 0,7 mm^2^, > 0,7 mm^2^) were selected at random for data collection (Suppl. Fig. 1). A previously published imaging mass cytometry panel developed for non-human primate (NHP) tissue was used for staining of markers of interest (20). Imaging revealed significant heterogeneity in granuloma organization, three granulomas representing this heterogeneity are depicted in figure 1D. T cell (middle column) localization differed strongly between granulomas; T cells are concentrated around the outer ring of the granuloma in the top granuloma, scattered throughout the lesion in the middle panel and directly surrounding the inner core of the lesion in the bottom panel of Figure 1D. Myeloid cell distribution also showed striking differences, illustrated in the right column of Figure 1D, with variable distribution of CD68, CD14 and myeloperoxidase expression and co-expression patterns indicated by yellow (CD68 and CD14), purple (CD14 and myeloperoxidase) and light blue (CD68 and myeloperoxidase). In the top and bottom granulomas, myeloperoxidase expressing neutrophils are confined to the core, while in the middle granuloma they are not found in a cluster but rather dispersed throughout the granuloma (right column, figure 1D). In the top granuloma CD14^+^ cells are found mostly in the outer ring, while they are more confined to the inner ring in the bottom granuloma. In both the top and bottom granulomas, there are clusters of CD68^+^ cell near the CD14^+^ cells, but while they are located in the inner ring in the top right granuloma, they are found in bigger clusters within the monocyte area in the bottom right granuloma (Figure 1D).

**Figure 1.**
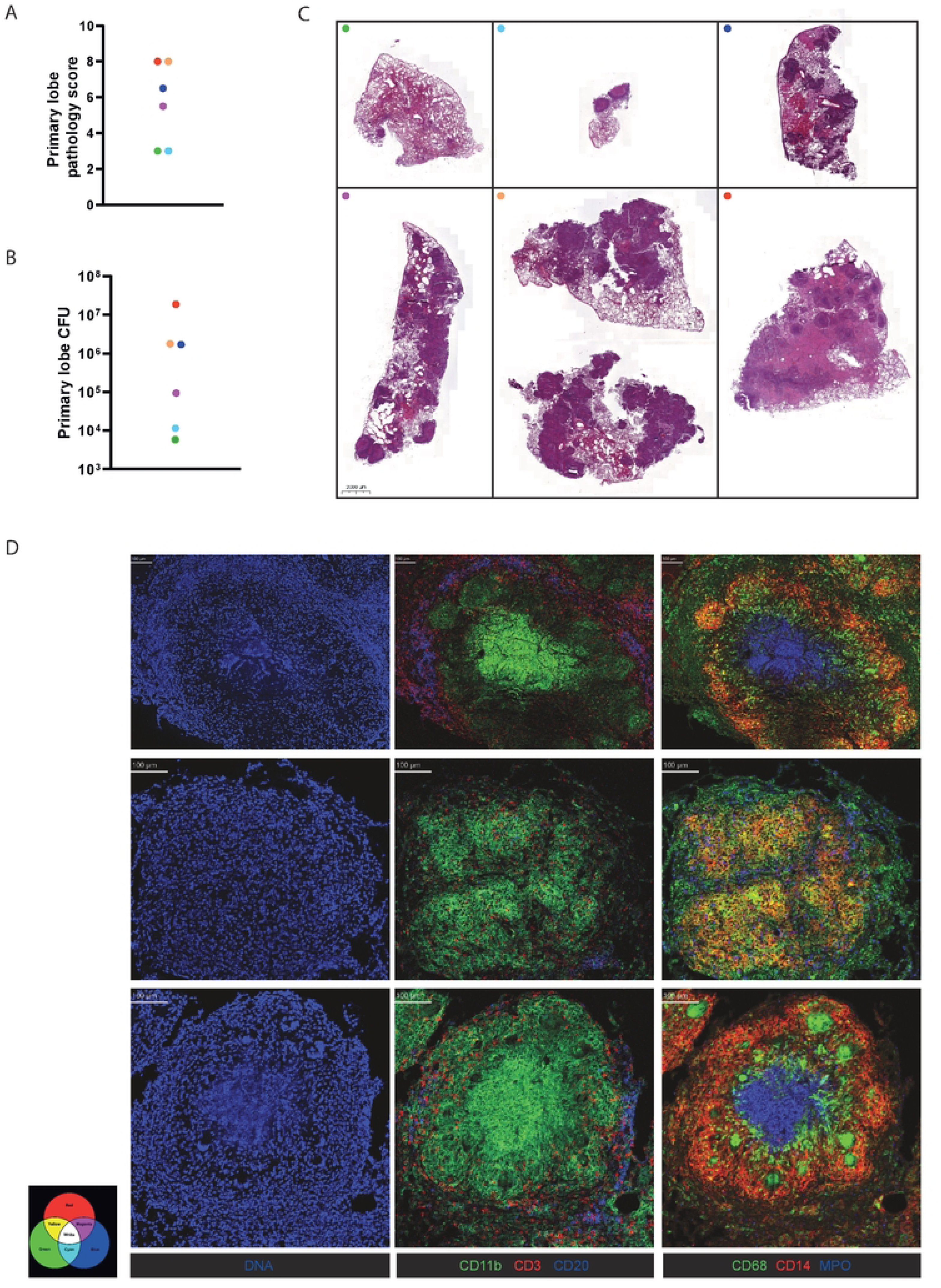
Disease parameters and lung images of *Mtb* Erdmann-infected macaques. (A+B) Lung pathology scores and primary lung lobe CFU burden of Mtb-infected macaques. (C) Hematoxylin and eosin staining of formalin fixed paraffin embedded lung tissue used in this study. (D) Visualization of 3 granuloma representative of the heterogeneity in the 28 imaged granuloma shown in 3 rows with the columns showing expression of different markers in these granuloma: (left column) DNA (blue), (middle column) CD11b+ myeloid cells (green), CD3+ T cells (red) and CD20+ B cells (blue) and (right column) CD68+ tissue macrophages (green), CD14+ monocytes (red) and myeloperoxidase+ neutrophils (blue).

To phenotype cells based on the expression of all markers measured, cell segmentation was performed to build single cell masks incorporating all pixels belonging to one cell into a single object. This creates what is effectively single cell data that can be clustered using t-Stochastic Neighbor Embedding (tSNE). Representative cell cluster images from all 28 investigated granulomas and associated heatmaps of marker expression are shown in Figure 2A and B, respectively. We identified calprotectin^+^ myeloperoxidase^+^ CD11b^+^ neutrophils, CD14^+^/CD68^+^ monocytes/macrophages, CD3^+^ T cells, CD3^-^CD7^+^ NK cells, CD20^+^ B cells, CD3^+^ TCRγδ^+^ T cells, CD117^+^ ILC, cytokeratin^+^ epithelial cells and undefined cells with heterogeneous distribution across all 28 granulomas (Figure 2C). Variation of cellular distributions between the granulomas was obvious for neutrophils, B cells and NK cells, while the macrophage population appeared relatively consistent. To identify factors of TB infection and disease associated with cellular granuloma composition, a principal component analysis was performed on the granuloma composition data and overlayed with other variables regarding animal species, granuloma characteristics, bacterial burden and gross pathology (Figure 2D). Variables that did not explain the variation in the cellular makeup were: macaque species, CFU burden of the lung, granuloma size and granuloma organization. However, when ranging granulomas by the individual animals’ severity of lung pathology, they clearly segregated in the PCA plot based on cellular composition (Figure 2D). Grouping cellular composition data based on this individual severity of lung disease revealed that granulomas from animals with limited lung pathology are enriched for NK cells, while intermediate pathology was associated an increased lymphoid compartment with highest proportions of T and B cells (Figure 2E). No significant differences were observed in the proportion of epithelial cells, macrophages, γδ T cells or ILC between these groups (Figure 2E and Suppl Fig. A-D).

**Figure 2.**
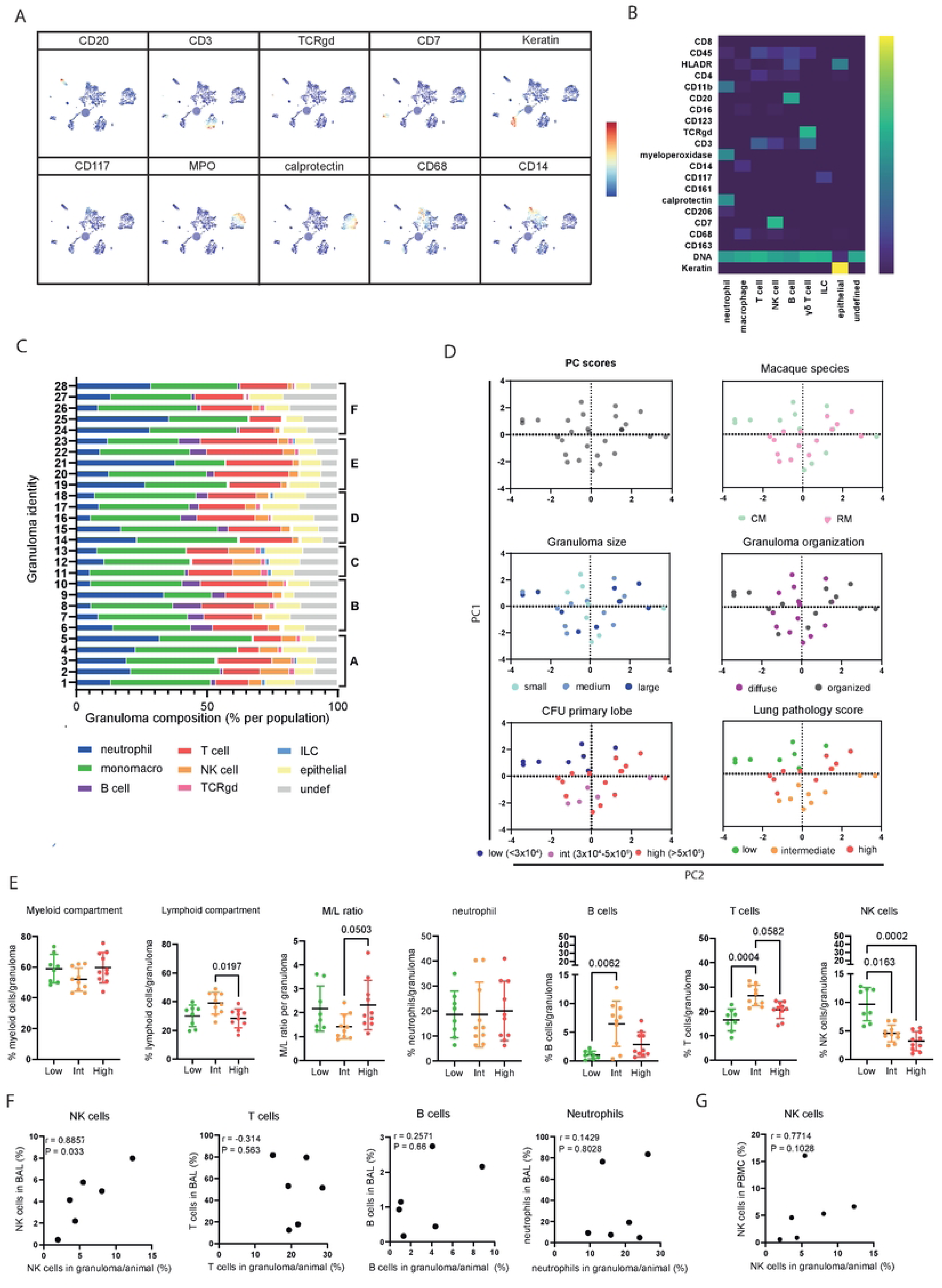
Granuloma composition correlates with lung pathology score and low pathology granuloma have increased proportions of NK cells. (A+B) Visualization of tSNE clustering of cells of all granuloma overlayed with expression density of several markers and heatmap of marker expression in different cell clusters. (C) Proportion of all identified populations in 28 granuloma. Letters refer to animal each imaged granuloma derives from. (D) PCA of granuloma according to cell population proportions (left top panel), overlayed with different parameters (macaque species, granuloma size, granuloma organization, CFU primary lobe and lung pathology score of the animals the granuloma were derived from). (E) Populations depicted as percentage of total cells in granuloma, grouped according to lung pathology scores low (green), intermediate (orange) and high (red). (F) Correlation plots of relationship between the average proportion of a cell type in the granuloma of an animal and the proportion of the same cell type in the bronchoalveolar lavage of the same animal at endpoint (6 weeks post-IGRA conversion). (G) Correlation plot of relationship between the average proportion of NK cells in the granuloma of an animal and the proportion of the same cell type in the PBMC of the same animal at baseline prior to infection.

As the similarity in composition of granulomas within animals (Figure 2C) and amongst animals with similar pathology levels (Figure 2E) suggests an animal-specific rather than a granuloma-specific composition signature, we hypothesized that such animal-specific variation in cell prevalence may also be reflected in other compartments. To test this hypothesis, the flow cytometry data on bronchoalveolar lavage cells (BALC) and peripheral blood mononuclear cells (PBMC) at 2 weeks prior to infection and 6 weeks post-IGRA conversion from the original Dijkman *et al.* NHP study were assessed to establish specific cell type frequencies (gating strategy in Suppl Fig. 3) (18). There was a significant positive correlation between the frequency of NK cells in the granulomas and in the BAL at endpoint (6 weeks post IGRA conversion) but not for any of the other subsets at these timepoints (Figure 2F, supplementary figure 2E). NK cell frequencies in PBMC prior to infection showed a trend of correlation with the NK cell frequency in granulomas at endpoint (Figure 2G). This suggests that NK cell abundance in granulomas is reflected in the local and systemic immune environment and not merely defined by the unique inflammatory milieu of each separate granuloma.

While there were no differences in the abundance of certain cell types between pathology groups, other parameters such as cellular interactions may differ. The preservation of spatial information in IMC data enables analysis of cellular interaction neighborhoods within the tissue. The cellular phenotypes identified above were overlayed onto the granuloma images (Figure 3A). Interaction analysis, based on cell proximity, was performed per individual lung pathology score group with each containing 8-10 granulomas. Results are shown in a grid, with the color of the box indicating the abundance of a particular interaction as percentage of the cell types’ total interactions, with blue as lowest number of interactions and yellow as highest proportion of interactions (Figure 3B). The grey and black dots indicate statistical significance of interactions, based on a permutation score with a cut-off of Z > 1.96 (equals p<0.05) (Figure 3B). In all granuloma categories, cell-cell interactions between cells of the same phenotype (e.g. neutrophils with neutrophils, T cells with T cells) were most common as expected. By comparing the significant interactions in the primary, low to high pathology score groups, interactions that were unique to each group were identified and indicated by bold black bordering of the respective cells in the grid (Figure 3B). Interestingly, NK cell-macrophage interactions were enriched in low pathology granulomas, suggesting this interaction may contribute to protective responses. Granulomas from animals with intermediate lung pathology showed more epithelial cell interactions with T and NK cells, and high pathology granulomas had increased γδ T cell interactions with both conventional T cells and NK cells. The increased interaction between NK cells and macrophages in low pathology lung granulomas is further highlighted by the images in Figure 3C, which only show cells involved in these interactions for one representative granuloma from each category. This finding is further represented by the relatively large proportion of NK cells in the total pool of macrophage interactions (Figure 3D) and of NK-macrophage microenvironments in the total pool of cellular interactions within granulomas (Figure 3E).

**Figure 3.**
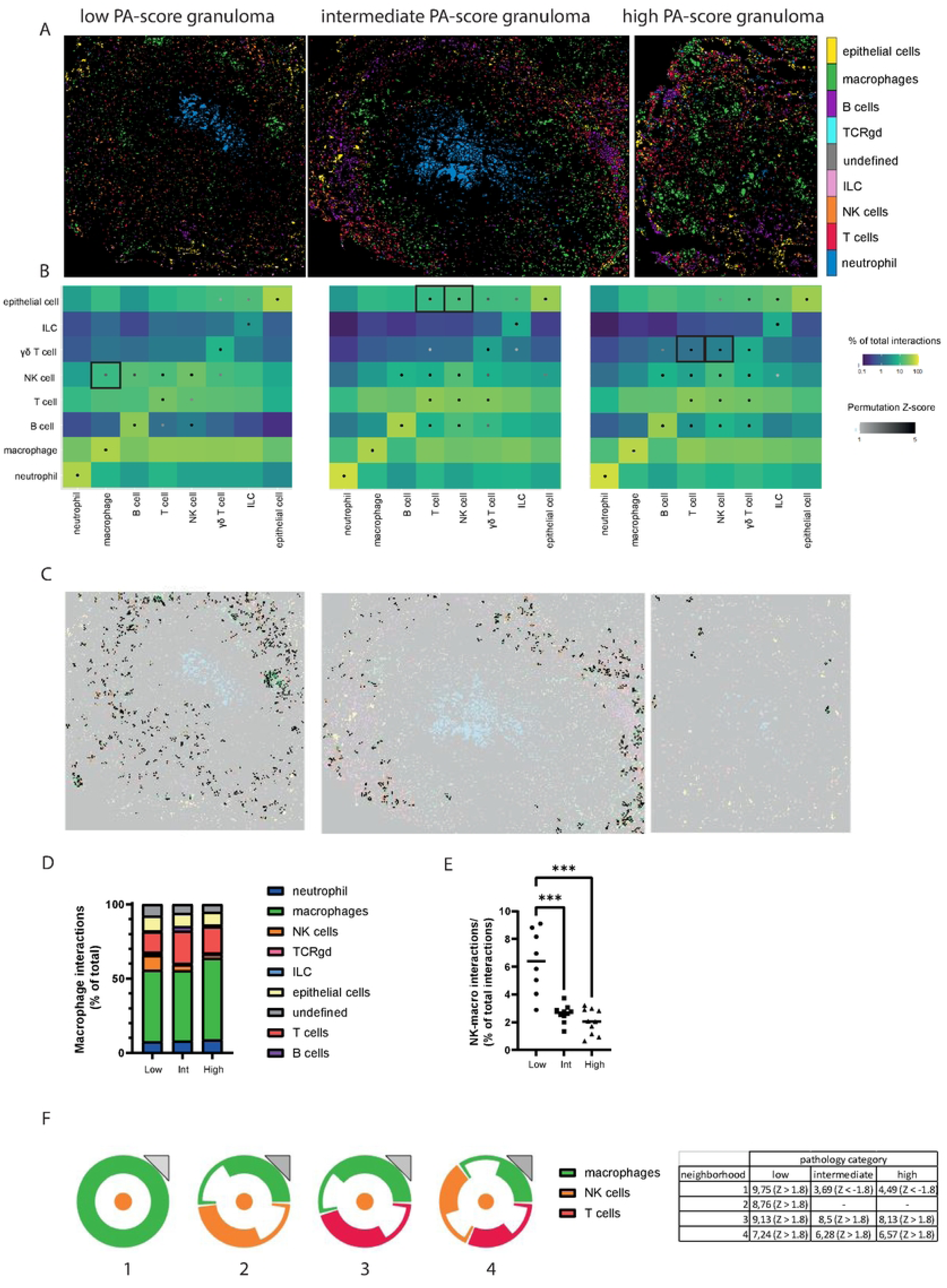
Granuloma from low pathology animals are enriched for NK cell-macrophage interactions. (A) Imaging mass cytometry images overlaid with cellular identities determined in figure 2A&B, showing one granuloma of each lung pathology score category (low, intermediate, high). (B) Interaction analysis for the granuloma of each lung pathology category with the colour of the square representing the frequency of the interaction between the phenotype of interest and the phenotype of neighborhood cells as a percentage of the total interactions for the phenotype of interest (blue to green gradient). The statistical analysis of the significant occurrence of an interaction is represented as a dot on the interaction square, showing only positive correlations (grey to black gradient). (C) Localization and frequency of NK cell-macrophage interactions in the three granuloma shown in figure A. (D) Frequency of cell interactions as frequency of total macrophage interactions. (E) Frequency of macrophage-NK cell (interactions in each granuloma as percentage of total interactions in the granuloma, shown across the lung pathology score categories. (F) Interaction glyphs showing an abstract representation of the four most abundant neighborhoods found in the macrophage-NK cell interactions and a table showing the proportion this neighborhood makes up of the macrophage-NK cell interactions, with the associated Z-score in brackets.

To identify other cell type interactions associated with the NK cell-macrophage microenvironment, interaction glyphs were created to visualize these neighborhoods in an abstracted image (Figure 3F). Four different neighborhoods were identified above the threshold of 50 interactions per granuloma; 1) an NK cell with surrounding macrophages, 2) an NK cell surrounded by an equal number of NK cells and macrophages, 3) an NK cell adjoined by an equal number of T cells and macrophages, 4) an NK cell neighbored by NK cells, macrophages and T cells. In the table the neighborhoods are shown as percentage of NK cell-macrophage interactions, which reveals similar percentages for neighborhoods 3 and 4 across pathology categories, while neighborhoods 1 and 2 with only NK cells and macrophages were mostly found in the granulomas of low pathology animals. Interaction 1 had a negative Z-score value for the granulomas from intermediate and high pathology score tissue, which indicates that finding NK cells with only surrounding macrophages occurs significantly less frequently than expected by chance on the basis the abundance of these cells.

As observed in figure 1, it is clear that there are multiple macrophage subsets in the granulomas. Therefore, we questioned whether a specific macrophage phenotype was selectively involved in the NK cell interactions in the primary lung lobe with limited pathology. Sub-clustering identified 4 macrophage subsets, based on expression levels of CD11b, CD14, CD16, CD68 and HLADR (Figure 4A and B); CD11b^+^ CD14^+^ CD16^int^ CD68^hi^ HLADR^+^ (designated Mac1), CD68^+^ CD14^lo^ (Mac2), CD14^+^ CD68^lo^ (Mac3), and CD16^int^ (Mac4). Neutrophils were also separated into 4 subclusters based on expression of CD11b, myeloperoxidase, calprotectin and CD206 as follows: CD11b^+^ MPO^hi^ calprotectin^+^ CD206^lo^ (designated Neutro1), CD11b^+^ MPO^hi^ (Neutro2), calprotectin^+^ (Neutro3), and CD11b^lo^ MPO^lo^ (Neutro4). The frequency of these macrophage and neutrophil subsets did not differ between granulomas by severity of lung pathology, with the exception of the Mac3 macrophage subset, which was increased in high pathology granulomas compared to low and intermediate pathology granulomas (Figure 4C and D). However, interaction analyses showed greater differences. Perhaps most notably, the majority of NK-macrophage interactions in low pathology granulomas was with Mac1 and Mac2, which are the CD68^+^ macrophage subsets (Figure 4E and F, respectively). While NK-Mac3 interactions did not differ between granuloma from different groups, NK-Mac4 interactions occurred more frequently in the high pathology group granulomas (Figure 4G-H). Next, we investigated which interactions were enriched in the distinct granuloma groups. In the intermediate and high pathology granulomas, interactions of both macrophage and neutrophil subsets are mostly with other macrophage and neutrophil subsets (Figure 4I, grey boxes), while only Mac4 showed significant interactions with other cell types, such as T and NK cells. In the granulomas from low pathology animals, the Mac1 subset shows interactions with NK cells, B cells and two of the neutrophil subsets (Figure 4I, black box). In all granulomas, regardless of level of pathology, macrophages belonging to subset 1 are found in clusters, while the other subsets appear scattered throughout the granulomas (Figure 4J). Although signals causing such clustering at this point remain elusive, it may hint to a relevant functional role for macrophage subset 1 interactions in association with local infection control. Taken together, spatial tissue analysis identifies intra-granuloma interactions associated with limited primary lung lobe pathology, pointing towards a relevant and preferential role of NK and CD11b^+^ CD14^+^ CD16^int^ CD68^hi^ HLADR^+^ macrophage interactions, contributing to our understanding of possible protective immune responses in the lung.

**Figure 4.**
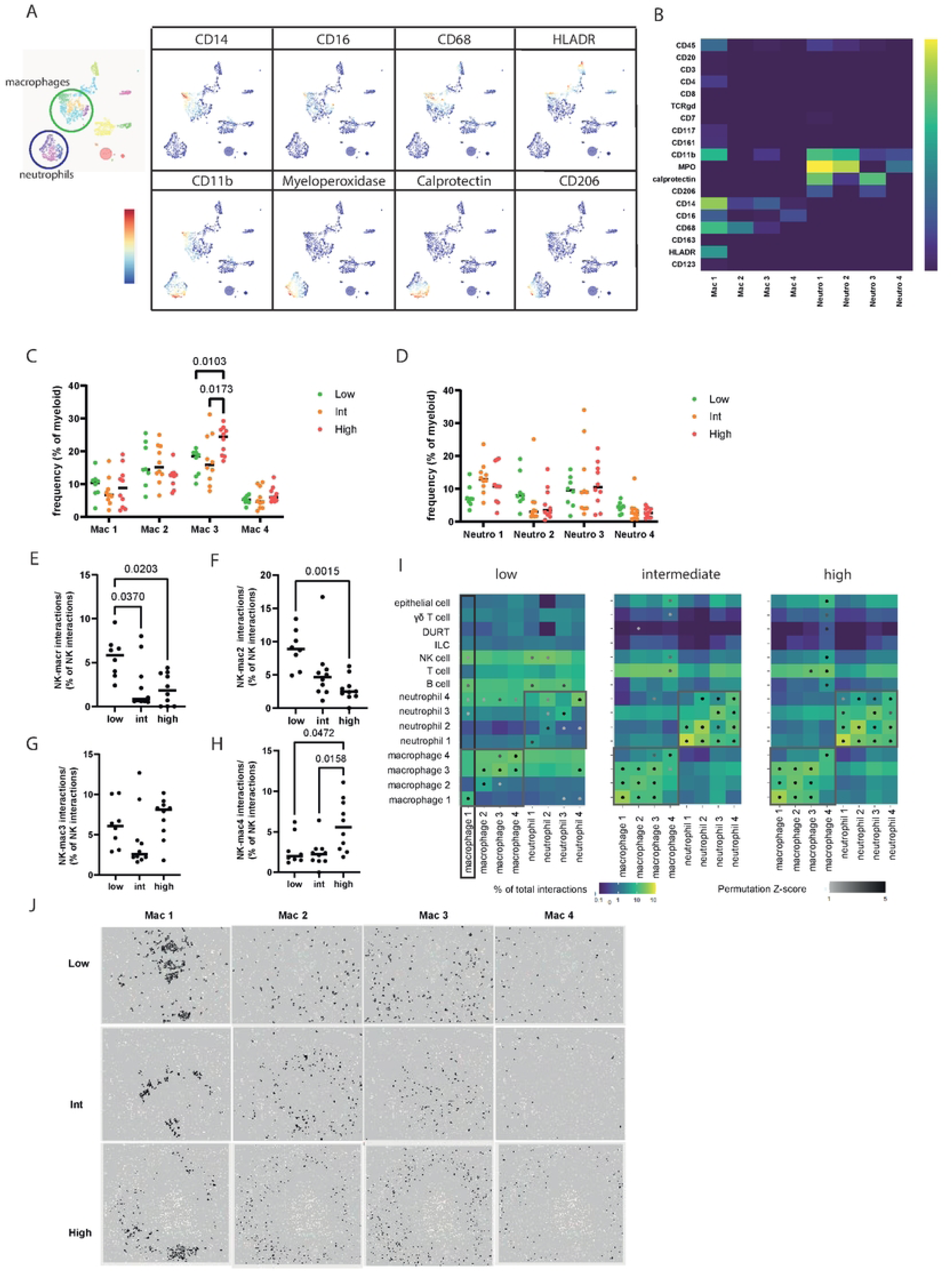
Detailed myeloid cell subtyping reveals tissue macrophage phenotype in NK cell interactions. (A-B) Visualization of clusters within the neutrophil (blue) and macrophage (green population) (A) and expression levels of relevant markers in map representation (A) and heatmap (B). (C-D) Distribution of macrophage (C) and neutrophil (D) subsets within the granuloma across lung pathology score groups (low = green, intermediate = orange, high = red). (E-H) Prevalence of NK cell interactions with the four different macrophage subsets in each granuloma across low, intermediate and high pathology animals, as percentage of total NK cell interactions. (I) Interaction analysis for the granuloma of each of the three lung pathology score groups (low, intermediate, high). The color of the square represents the frequency of the interaction between the phenotype of interest and the phenotype of neighborhood cells as a percentage of the total interactions for the phenotype of interest (blue to green gradient). The statistical analysis of the significant occurrence of an interaction is represented as a dot on the interaction square (grey to black gradient). The grey boxes indicate the cellular interactions within the neutrophil and macrophage lineage and the black box shows the interactions of macrophage 1 subset with different cell types in the low lung pathology score group. (J) Visualization of location and grouping of the four macrophage subsets in a representative granuloma of each of the pathology categories (low, intermediate and high).

## Discussion

The formation of granulomas at the site of infection is a hallmark of tuberculosis disease and these structures reflect the primary site of host-pathogen interactions, rendering them suitable to study immune responses against *Mtb*. Granulomas are notoriously heterogenous, even within the same individual (5, 21). Several studies have performed elegant analyses on individual granulomas to identify factors related to the mycobacterial burden within the micro-environment of an individual granulomas (22, 23). As outcome of disease likely comes down to overall capacity to control bacterial load, studying the relationship between granuloma composition and other parameters of disease may also be informative. Here, we show that despite their superficial heterogeneity in visual appearance and size, similarities between non-human primate granulomas are evident when categorized by the pathological involvement of the respective lung lobe from which the granuloma originate. Interestingly, granulomas from the same animal analyzed by imaging mass cytometry in this study, do show cell distributions that appear more similar than distinct, suggesting that granuloma composition may be a factor of the whole lung environment or even the systemic immune response, rather than the granuloma’s microenvironment. This notion is supported by the similarity of granuloma composition between different animals with comparable levels of lung pathology. Here, we identified immune signatures of specific cell interactions associated with limited severity of lung pathology relatively early in infection.

Assessing all immune subsets separately showed that both B and T cells are most abundant in granulomas of animals with intermediate lung pathology severity. The high number of B cells in intermediate pathology granulomas is hard to interpret as the role of B cells in anti-*Mtb* responses is unclear and their abundance does not seem directly correlated to protection, although numbers are reduced in infected individuals and they may have an indirect role in the context of vaccination (Joosten, 2016, Plos Pathogen; Chowdhury, 2018, Nature Letters, Slight, 2013, JCI). On the other hand, T cells are considered one of the most important cell types to provide protection against intracellular bacteria such as *Mtb* and increased recruitment of T cells is generally regarded as desirable for protection from disease. Gautam *et al*. reported that improved T cell positioning in the granuloma, achieved by depleting immunomodulatory IDO, resulted in improved bacterial killing (8) (24), but despite varying T cell proportions we did not observe altered T cell positioning between granulomas from animals with differing severity scores. The increased numbers of T cells may not impact pathology or bacterial load here, as it has been established that only a small percentage of T cells in the granuloma are *Mtb*-responsive and only T cells with a particular balance of pro- and anti-inflammatory cytokine production correlate with clearance of *Mtb* (25) (9). The contribution of Treg was not assessed separately as they could not be identified with the current panel and are therefore included in the total T cell count.

NK cells were most abundant in granulomas of animals with low pathology compared to intermediate and high pathology animal granulomas, suggesting these cells may be involved in controlling disease. This is in line with findings in several models of TB disease. Murine mucosal BCG vaccination resulted in expansion of tissue-resident NK cells, which contributed to protection against *Mtb* infection (26). A previous study in NHP using single cell RNA analyses also showed that (CD27^+^) NK cell abundance was increased in the lungs of animals with controlled disease compared to uncontrolled disease (11, 27). A potential protective role for NK cells is further supported by studies in human TB cohorts showing positive correlations between abundance of specific peripheral blood NK cell subsets and resistance to disease progression, while NK cell numbers are reduced during active disease (28–31).

Previously, a study in South African human cohorts identified a negative correlation between peripheral NK cell numbers and the level of inflammation in the lung, as measured by PET-CT scan (29). Here, we confirm the negative correlation between NK cell numbers and severity of lung pathology, and expand on this finding by showing that peripheral NK cell numbers are in concordance with those in the airways and granulomas. This further supports a potential protective role for NK cells in the tissue as well as a broader immune signature across the lung and suggests it may be possible to track certain immune events in the lung parenchyma peripherally. Indeed, previous work correlated immune (regulatory) signatures found in human granuloma with the blood transcriptome (15).

The preservation of spatial context in imaging mass cytometry data enables analysis of the cell-cell neighborhoods in the granulomas across the lung pathology gradient. Here, granulomas from animals with limited pathology showed significantly elevated interactions between NK cells and macrophage subsets with a CD68^+^ tissue-resident macrophage phenotype in comparison to granulomas from animals with more severe pathology. This suggests these cells and their interaction may be important for protective responses against *Mtb*. Mechanisms of action against *Mtb* may include direct killing of free *Mtb* by NK cells (32), killing of *Mtb*-infected macrophages by NK cells (33), cytokine-mediated stimulation of phagolysosomal fusion in *Mtb*-containing macrophages (34) or indirect activation of other immune subsets (35). In future studies, it would be of interest to analyze NK cell-tissue macrophages interactions in more detail, as these may be the cells most effective in collaborating to kill the mycobacteria. To our knowledge, this is the first study identifying NK cells and their interactions in the granuloma as possible contributors to mycobacterial control.

As a whole, this work shows detailed analysis of the cellular make-up and cell neighborhoods in non-human primate granulomas. It identifies a correlation between lung pathology score and granuloma composition and shows an increase in NK cell-tissue macrophage interactions in animals with relatively limited TB disease manifestation.

## Materials & methods

### Tissues

Archived NHP tissue samples were obtained from the Biomedical Primate Research Centre (BPRC), Rijswijk, the Netherlands (courtesy of DVM I. Kondova). BPRC is licensed by the Dutch authority to breed non-human primates and to use them for research in areas of life-threatening and disabling diseases without suitable alternatives. BPRC complies to all relevant legislation with regard to the use of animals in research; the Dutch ‘*Wet op de Dierproeven*’ and the European guideline 2010/63/EU. BPRC is AAALAC accredited since 2012. All TB samples used in this study were derived from the biobank of a previously performed study, registered under DEC accession no. 761subA (18). Thus, by their availability, samples were exploited beyond legal requirement for prior approval of protocol, while no additional animal study was performed for the purpose of the experiments described in this manuscript.

Here, we used formalin-fixed paraffin-embedded (FFPE) lung tissue that was collected from 3 rhesus and 3 cynomolgus macaques 6 weeks after a negative to positive conversion of an antigen-specific interferon-gamma response assay (IGRA; *in casu* a NHP IFNg-specific ELISPOT labtest) corroborating *Mtb* infection upon low dose *Mtb* Erdman challenge via endobronchial instillation (18). In addition, we further analyzed data from previously performed flow cytometry experiments on bronchoalveolar lavage cells (BALC) and peripheral blood mononuclear cells (PBMC) from the same animals (18). The primary endobronchially targeted lung lobe was scored for pathological involvement as previously described (Dijkman et al FIMMU 2019), yielding arbitrary units of one to four for both prevalence (1-3, 4-10, >10 lesions, or miliary/coalescing manifestation) and size of lesions (< 2 mm, 2-4 mm, > 4 mm, coalescing).

### Hematoxylin and eosin staining

4 µm sections were cut from FFPE blocks of lung tissue and transferred to silane-coated glass slides (VWR, Radnor, PA, USA). Slides were dried overnight and deparaffinized through an xylol and ethanol gradient. Next, slides were washed with MilliQ and stained with Mayer’s hematoxylin (Sigma-Aldrich, St. Louis, MO, USA) for 4 minutes. After a 50 minute wash under running tap water, the slides were stained with Eosin (Sigma-Aldrich, St. Louis, MO, USA) for 1 minute and subsequently dehydrated. Finally, mounting medium was applied and slides were cover slipped. A 3D Histech Pannoramic 250 Flash III slide scanner (3D Histech, Budapest, Hungary) was used to scan the slides at ×40 magnification.

### Imaging mass cytometry staining

Imaging mass cytometry staining was performed on 4 µm sections cut from FFPE blocks of NHP lung tissue as previously described (20, 36). Briefly, slides were dried in an oven at 37°C overnight and deparaffinized through sequential xylene and ethanol washes. Slides were then washed in 1x low pH antigen retrieval solution (Thermo Fisher Scientific) and subsequently boiled in pre-heated antigen retrieval solution for 10 minutes. After cooling, slides were washed with 1x PBS-TB (PBS containing 0.05% Tween-20 and 1% BSA) and incubated with Superblock solution (Thermo Fisher Scientific, Waltham, MA, USA) for 30 min at RT. Slides were incubated with 100 μL anti-CD8α antibody in PBS-TB overnight at 4°C in a humid chamber, washed several times with PBS-TB and then incubated with secondary antibody Dy800 for 1 h at RT the following day. Next, slides were washed several times and then incubated with 100 μL antibody mix containing all remaining antibodies overnight at 4°C (Table 1). After washing, slides were incubated with intercalator (Ir) in PBS-TB for 5 min at RT. Finally, slides were washed with PBS-TB and water and dried overnight.

**Table 1.**
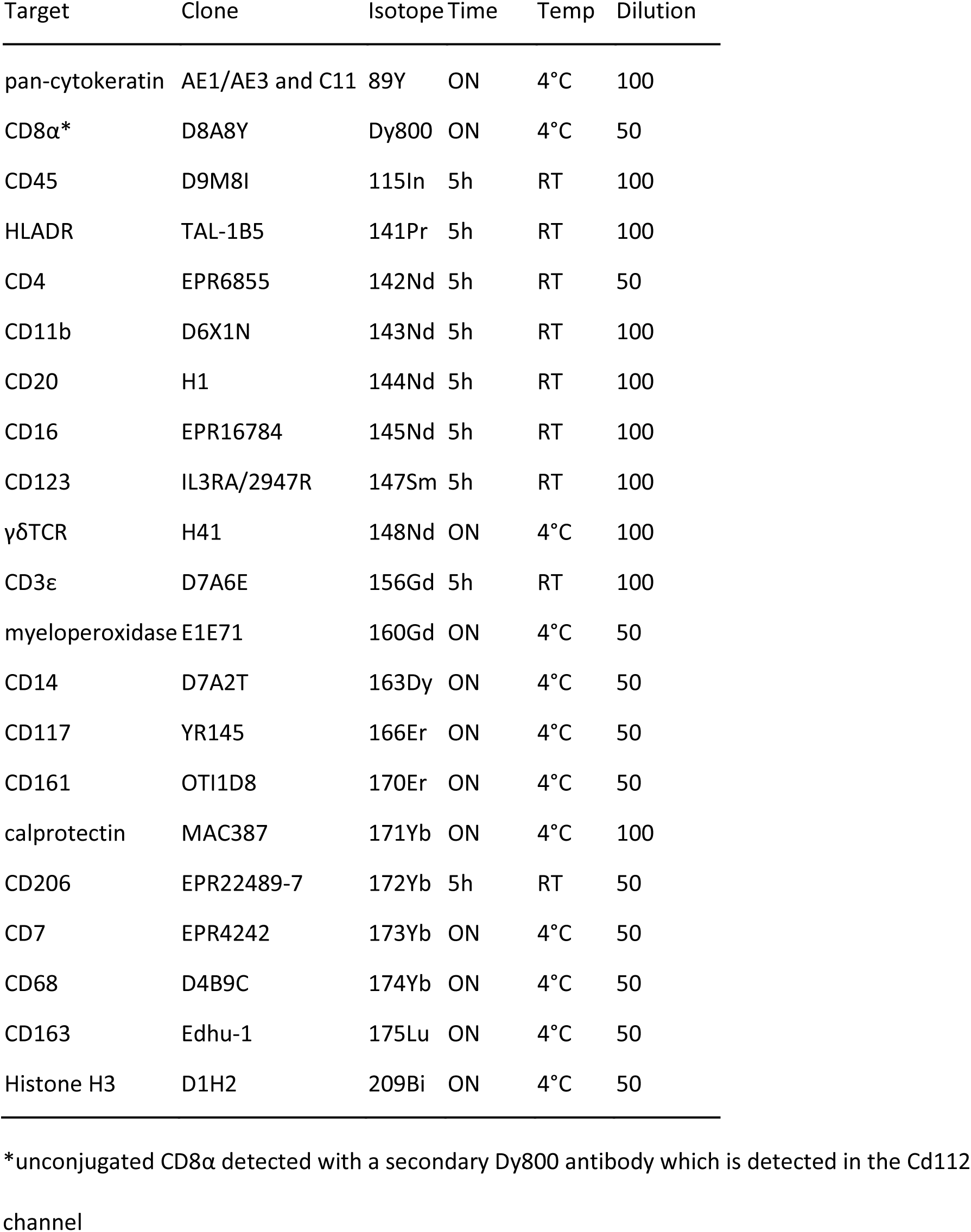
21-marker panel for imaging mass cytometry, with antibody-metal combination, antibody clone and staining conditions.

| Target | Clone | Isotope | Time | Temp | Dilution |
| --- | --- | --- | --- | --- | --- |
| pan-cytokeratin | AE1/AE3 and C11 | 89Y | ON | 4°C | 100 |
| CD8α* | D8A8Y | Dy800 | ON | 4°C | 50 |
| CD45 | D9M8I | 115In | 5h | RT | 100 |
| HLADR | TAL-1B5 | 141Pr | 5h | RT | 100 |
| CD4 | EPR6855 | 142Nd | 5h | RT | 50 |
| CD11b | D6X1N | 143Nd | 5h | RT | 100 |
| CD20 | H1 | 144Nd | 5h | RT | 100 |
| CD16 | EPR16784 | 145Nd | 5h | RT | 100 |
| CD123 | IL3RA/2947R | 147Sm | 5h | RT | 100 |
| γδTCR | H41 | 148Nd | ON | 4°C | 100 |
| CD3ε | D7A6E | 156Gd | 5h | RT | 100 |
| myeloperoxidase | E1E71 | 160Gd | ON | 4°C | 50 |
| CD14 | D7A2T | 163Dy | ON | 4°C | 50 |
| CD117 | YR145 | 166Er | ON | 4°C | 50 |
| CD161 | OTI1D8 | 170Er | ON | 4°C | 50 |
| calprotectin | MAC387 | 171Yb | ON | 4°C | 100 |
| CD206 | EPR22489-7 | 172Yb | 5h | RT | 50 |
| CD7 | EPR4242 | 173Yb | ON | 4°C | 50 |
| CD68 | D4B9C | 174Yb | ON | 4°C | 50 |
| CD163 | Edhu-1 | 175Lu | ON | 4°C | 50 |
| Histone H3 | D1H2 | 209Bi | ON | 4°C | 50 |

\*unconjugated CD8α detected with a secondary Dy800 antibody which is detected in the Cd112 channel

### Imaging mass cytometry acquisition

All mass cytometry data was acquired on the Hyperion mass cytometry imaging system (Standard Biotools, San Fransisco, CA, USA) at the Flow cytometry Core Facility at the LUMC. A tuning protocol and 3-element tuning slide were used to autotune the system, as per the Hyperion imaging system user guide (Standard Biotools). Areas of interest were identified on consecutive H&E slides by light microscopy. Selected areas were ablated and acquired at 200 Hz, after which data was exported as MCD files. These files were visualized using the MCD viewer.

### Imaging mass cytometry data analysis

Data normalization was performed in Ilastik by using semi-automated background removal to control for signal-to-noise variations between FFPE sections (37, 38). Cell segmentation masks were created in ilastik and cell profiler (39) and combined with normalized images in ImaCyte to generate FCS files containing single cell data on positive pixel frequency for each marker. Cytosplore was used to perform hierarchical Stochastic Neighbour Embedding (hSNE) with 1000 iterations and 3 scales (40). The resulting cell clusters were exported as FCS files. Segmentation masks and the FCS subsets were loaded into Imacyte to enable spatial visualization and perform microenvironment analysis (41). The settings used for this analysis were a distance of 5 pixels and a minimum of 20 occurrences of a neighborhood. A permutation Z-score was calculated for each interaction and a cutoff of 1.96 (p < 0.05) was used. Frequency of occurrence of microenvironments was calculated based on the count of at least one co-localized cell divided by the total number of interactions or the number of cells in the cluster of that sample.

### Flow cytometry data

The flow cytometry data shown in this paper was collected as reported by Dijkman *et al.*(18) and reanalyzed in FlowJo (Treestar Inc, Ashland, OR, USA) to obtain frequencies of populations of interest in PBMC and BALC.

## Data availability

The imaging mass cytometry data are available at https://figshare.com/s/efba49c2eee333c15b35

## Author contributions

Study design: PN, TO, FV, SJ

Conducting experiments & acquiring data: PN, KD

Analyzing data: PN, MIJ, FV

Writing the manuscript: PN

Editing the manuscript: MIJ KD NM TO FV SJ

## Acknowledgements

The authors would like to acknowledge the contribution of Richard Vervenne from the BPRC for the assessment of the flow cytometry data and DVM I. Kondova in contributing to the assessment providing the tissue blocks from the BPRC.

## Notes

### Competing Interest Statement

The authors have declared no competing interest.

## References

1. WHO. Global tuberculosis report 2024. Geneva 2024.

2. Palomino JC, Martin A. Drug Resistance Mechanisms in Mycobacterium tuberculosis. Antibiotics (Basel). 2014;3(3):317–40.

3. Kumar P. A Perspective on the Success and Failure of BCG. Front Immunol. 2021;12:778028.

4. Pai M, Behr MA, Dowdy D, Dheda K, Divangahi M, Boehme CC, et al. Tuberculosis. Nat Rev Dis Primers. 2016;2:16076.

5. Lin PL, Ford CB, Coleman MT, Myers AJ, Gawande R, Ioerger T, et al. Sterilization of granulomas is common in active and latent tuberculosis despite within-host variability in bacterial killing. Nat Med. 2014;20(1):75–9.

6. Carow B, Hauling T, Qian X, Kramnik I, Nilsson M, Rottenberg ME. Spatial and temporal localization of immune transcripts defines hallmarks and diversity in the tuberculosis granuloma. Nat Commun. 2019;10(1):1823.

7. Cronan MR. In the Thick of It: Formation of the Tuberculous Granuloma and Its Effects on Host and Therapeutic Responses. Front Immunol. 2022;13:820134.

8. Gautam US, Foreman TW, Bucsan AN, Veatch AV, Alvarez X, Adekambi T, et al. In vivo inhibition of tryptophan catabolism reorganizes the tuberculoma and augments immune-mediated control of Mycobacterium tuberculosis. Proc Natl Acad Sci U S A. 2018;115(1):E62–E71.

9. Millar JA, Butler JR, Evans S, Mattila JT, Linderman JJ, Flynn JL, et al. Spatial Organization and Recruitment of Non-Specific T Cells May Limit T Cell-Macrophage Interactions Within Mycobacterium tuberculosis Granulomas. Front Immunol. 2020;11:613638.

10. Ndlovu H, Marakalala MJ. Granulomas and Inflammation: Host-Directed Therapies for Tuberculosis. Front Immunol. 2016;7:434.

11. Esaulova E, Das S, Singh DK, Choreno-Parra JA, Swain A, Arthur L, et al. The immune landscape in tuberculosis reveals populations linked to disease and latency. Cell Host Microbe. 2021;29(2):165–78 e8.

12. Jackson HW, Fischer JR, Zanotelli VRT, Ali HR, Mechera R, Soysal SD, et al. The single-cell pathology landscape of breast cancer. Nature. 2020;578(7796):615–20.

13. Schwabenland M, Salie H, Tanevski J, Killmer S, Lago MS, Schlaak AE, et al. Deep spatial profiling of human COVID-19 brains reveals neuroinflammation with distinct microanatomical microglia-T-cell interactions. Immunity. 2021;54(7):1594–610 e11.

14. van Unen V, Ouboter LF, Li N, Schreurs M, Abdelaal T, Kooy-Winkelaar Y, et al. Identification of a Disease-Associated Network of Intestinal Immune Cells in Treatment-Naive Inflammatory Bowel Disease. Front Immunol. 2022;13:893803.

15. McCaffrey EF, Donato M, Keren L, Chen Z, Delmastro A, Fitzpatrick MB, et al. The immunoregulatory landscape of human tuberculosis granulomas. Nat Immunol. 2022;23(2):318–29.

16. Sawyer AJ, Patrick E, Edwards J, Wilmott JS, Fielder T, Yang Q, et al. Spatial mapping reveals granuloma diversity and histopathological superstructure in human tuberculosis. J Exp Med. 2023;220(6).

17. Scanga CA, Flynn JL. Modeling tuberculosis in nonhuman primates. Cold Spring Harb Perspect Med. 2014;4(12):a018564.

18. Dijkman K, Vervenne RAW, Sombroek CC, Boot C, Hofman SO, van Meijgaarden KE, et al. Disparate Tuberculosis Disease Development in Macaque Species Is Associated With Innate Immunity. Front Immunol. 2019;10:2479.

19. Giesen C, Wang HA, Schapiro D, Zivanovic N, Jacobs A, Hattendorf B, et al. Highly multiplexed imaging of tumor tissues with subcellular resolution by mass cytometry. Nat Methods. 2014;11(4):417–22.

20. Niewold P, Ijsselsteijn ME, Verreck FAW, Ottenhoff THM, Joosten SA. An imaging mass cytometry immunophenotyping panel for non-human primate tissues. Front Immunol. 2022;13:915157.

21. Lenaerts A, Barry CE, 3rd, Dartois V. Heterogeneity in tuberculosis pathology, microenvironments and therapeutic responses. Immunol Rev. 2015;264(1):288–307.

22. Gideon HP, Hughes TK, Tzouanas CN, Wadsworth MH, Tu AA, Gierahn TM, et al. Multimodal profiling of lung granulomas in macaques reveals cellular correlates of tuberculosis control. Immunity. 2022;55(5):827-+.

23. Marakalala MJ, Raju RM, Sharma K, Zhang YJJ, Eugenin EA, Prideaux B, et al. Inflammatory signaling in human tuberculosis granulomas is spatially organized. Nature Medicine. 2016;22(5):531–8.

24. Singh B, Moodley C, Singh DK, Escobedo RA, Sharan R, Arora G, et al. Inhibition of indoleamine dioxygenase leads to better control of tuberculosis adjunctive to chemotherapy. JCI Insight. 2023;8(2).

25. Gideon HP, Phuah J, Myers AJ, Bryson BD, Rodgers MA, Coleman MT, et al. Variability in Tuberculosis Granuloma T Cell Responses Exists, but a Balance of Pro- and Anti-inflammatory Cytokines Is Associated with Sterilization. Plos Pathogens. 2015;11(1).

26. Durojaye O, Vankayalapati A, Paidipally P, Mukherjee T, Vankayalapati R, Radhakrishnan RK. Lung-resident CD3-NK1.1+CD69+CD103+ Cells Play an Important Role in Bacillus Calmette-Guerin Vaccine-Induced Protective Immunity against Mycobacterium tuberculosis Infection. J Immunol. 2024;213(5):669–77.

27. Darrah PA, Zeppa JJ, Wang C, Irvine EB, Bucsan AN, Rodgers MA, et al. Airway T cells are a correlate of i.v. Bacille Calmette-Guerin-mediated protection against tuberculosis in rhesus macaques. Cell Host Microbe. 2023;31(6):962–77 e8.

28. Cai Y, Dai Y, Wang Y, Yang Q, Guo J, Wei C, et al. Single-cell transcriptomics of blood reveals a natural killer cell subset depletion in tuberculosis. EBioMedicine. 2020;53:102686.

29. Chowdhury RR, Vallania F, Yang QT, Angel CJL, Darboe F, Penn-Nicholson A, et al. A multi-cohort study of the immune factors associated with infection outcomes. Nature. 2018;560(7720):644-+.

30. Venkatasubramanian S, Cheekatla S, Paidipally P, Tripathi D, Welch E, Tvinnereim AR, et al. IL-21-dependent expansion of memory-like NK cells enhances protective immune responses against Mycobacterium tuberculosis. Mucosal Immunol. 2017;10(4):1031–42.

31. Zhou Y, Lan H, Shi H, Wu P, Zhou Y. Evaluating the diversity of circulating natural killer cells between active tuberculosis and latent tuberculosis infection. Tuberculosis (Edinb). 2022;135:102221.

32. Esin S, Batoni G, Pardini M, Favilli F, Bottai D, Maisetta G, et al. Functional characterization of human natural killer cells responding to Mycobacterium bovis bacille Calmette-Guerin. Immunology. 2004;112(1):143–52.

33. Brill KJ, Li Q, Larkin R, Canaday DH, Kaplan DR, Boom WH, et al. Human natural killer cells mediate killing of intracellular Mycobacterium tuberculosis H37Rv via granule-independent mechanisms. Infect Immun. 2001;69(3):1755–65.

34. Dhiman R, Indramohan M, Barnes PF, Nayak RC, Paidipally P, Rao LV, et al. IL-22 produced by human NK cells inhibits growth of Mycobacterium tuberculosis by enhancing phagolysosomal fusion. J Immunol. 2009;183(10):6639–45.

35. Vankayalapati R, Klucar P, Wizel B, Weis SE, Samten B, Safi H, et al. NK cells regulate CD8+ T cell effector function in response to an intracellular pathogen. J Immunol. 2004;172(1):130–7.

36. Ijsselsteijn ME, van der Breggen R, Farina Sarasqueta A, Koning F, de Miranda N. A 40-Marker Panel for High Dimensional Characterization of Cancer Immune Microenvironments by Imaging Mass Cytometry. Front Immunol. 2019;10:2534.

37. Berg S, Kutra D, Kroeger T, Straehle CN, Kausler BX, Haubold C, et al. ilastik: interactive machine learning for (bio)image analysis. Nat Methods. 2019;16(12):1226–32.

38. Ijsselsteijn ME, Somarakis A, Lelieveldt BPF, Hollt T, de Miranda N. Semi-automated background removal limits data loss and normalizes imaging mass cytometry data. Cytometry A. 2021;99(12):1187–97.

39. Carpenter AE, Jones TR, Lamprecht MR, Clarke C, Kang IH, Friman O, et al. CellProfiler: image analysis software for identifying and quantifying cell phenotypes. Genome Biol. 2006;7(10):R100.

40. Höllt T, Pezzotti N, van Unen V, Koning F, Eisemann E, Lelieveldt B, et al. Cytosplore: Interactive Immune Cell Phenotyping for Large Single-Cell Datasets. Comput Graph Forum. 2016;35(3):171–80.

41. Somarakis A, Van Unen V, Koning F, Lelieveldt B, Hollt T. ImaCytE: Visual Exploration of Cellular Micro-Environments for Imaging Mass Cytometry Data. IEEE Trans Vis Comput Graph. 2021;27(1):98–110.

